# OHNOLOGS v2: A comprehensive resource for the genes retained from whole genome duplication in vertebrates

**DOI:** 10.1101/717124

**Authors:** Param Priya Singh, Hervé Isambert

## Abstract

All vertebrates including human have evolved from an ancestor that underwent two rounds of whole genome duplication (2R-WGD). In addition, teleost fish underwent an additional third round of genome duplication (3R-WGD). The genes retained from these genome duplications, so-called ohnologs, have been instrumental in the evolution of vertebrate complexity, developmental patterns and susceptibility to genetic diseases. However, the identification of vertebrate ohnologs has been challenging, due to lineage specific genome rearrangements since 2R- and 3R-WGD. We have previously identified vertebrate ohnologs using a novel synteny comparison across multiple genomes. Here, we refine and apply this approach on 27 vertebrate genomes to identify ohnologs from both 2R- and 3R-WGD, while taking into account the phylogenetically biased sampling of available species. We assemble vertebrate ohnolog pairs and families in an expanded OHNOLOGS v2 database, which also includes non-protein coding RNA genes. We find that teleost fish have retained most 2R-WGD ohnologs common to amniotes, which have also retained significantly more ohnologs from 3R-WGD, whereas a higher rate of 2R-WGD ohnolog loss is observed in sauropsids compared to mammals and fish. OHNOLOGS v2 should allow deeper evolutionary genomic analysis of the impact of WGD on vertebrates and can be freely accessed at http://ohnologs.curie.fr.

## INTRODUCTION

Gene duplication provides raw material for evolution of new gene functions (1). Duplication of single genes or genomic segments is a continuous evolutionary process that creates diversity in terms of copy number variations (CNV) across individuals, and paralogs across species. In addition, dramatic evolutionary accidents corresponding to Whole Genome Duplication (WGD) have also occurred in the evolutionary past of most eukaryotic lineages including plans, fungi, and animals (2–4). For example, all extant vertebrates have experienced two rounds of whole genome duplications (2R-WGD) in their evolutionary past (5–8). In addition, a third round of genome duplication have also occurred in the teleost fish lineage (3R-WGD) (9–11). 2R-WGDs likely played important role in the evolution and diversification of vertebrate specific innovations such as neural crest cells, placodes, and a complex brain (12,13). Many key genes implicated in development of these structures can be traced back to 2R-WGD. Similarly, 3R-WGD likely played an important role in the expansion of the diversity of teleost fish lineage making it the most species rich vertebrate group (14–17). Hence, the genes retained from these three WGD events have been instrumental in the evolution of vertebrates (18).

The genes originated from these ancient polyploidy (paleo-polyploidy) events are now called ohnologs after Susumu Ohno who first hypothesized the two rounds of WGD events in vertebrate ancestors (1,5,19). Ohnologs are known to have distinct evolutionary, genomic and functional properties that distinguish them from small-scale duplicates and singletons (20–23). They also show greater association with diseases and cancer than non-ohnolog genes (24–29), and have been suggested to be dosage balanced (24), which was subsequently shown to be indirectly mediated by their high susceptibility to dominant mutations (25,27,28).

Given the specific impact WGDs have had on the evolution of vertebrates, a comprehensive database of vertebrate ohnologs is highly desirable. While there are some useful resources available for comparison of synteny across species (30–33) there is no database that reliably identifies ohnologs from both vertebrate 2R-WGDs and fish 3R-WGD. To start filling this gap, we developed in 2015, OHNOLOGS, a repository of ohnologs retained from the 2R-WGD in six amniote vertebrates (human, mouse rat, pig, dog and chicken) (33). OHNOLOGS is based on a novel comparative macro-synteny approach that reliably identifies ohnologs, despite lineage specific genome rearrangement, gene loss and small scale duplication events, by combining macro-synteny information (gene content regardless of exact order) across multiple outgroups and vertebrate genomes (33).

Here, we expand this multiple genome synteny comparison approach to 27 vertebrate species including 4 teleost fish species. We further improve the statistical confidence assessment of each ohnolog pair with a weighted quantitative confidence score (q-score) taking into account the phylogenetically biased sampling of available vertebrate species. In addition, we uncover ohnologs, including in non-protein coding RNA gene classes, from both 2R-WGD in early vertebrates (2R-ohnologs) and 3R-WGD in teleost fish (3R-ohnologs). The expanded OHNOLOGS database is the most comprehensive repository of ohnologs in vertebrates. Using the new OHNOLOGS database we show that the average 2R- and 3R-ohnolog retention rates are 25% and 18% respectively, with sauropsida showing the highest lineage specific loss of 2R-ohnologs, and teleost fish showing the highest lineage specific retention of 2R-ohnologs. We found that 2R-ohnologs are significantly more likely to also retain 3R-ohnologs, in agreement with earlier reports (34). OHNOLOGS v2 should facilitate deeper evolutionary analysis of the unique properties of ohnologs, and their lineage specific retention and loss in different vertebrates.

## RESULTS

### Data collection and processing

OHNOLOGS v2 includes 2R-ohnolog pairs and families in 27 vertebrates that have a chromosome level assembly with a majority of their genes anchored on chromosomes in Ensembl version 84 (35). This includes 18 mammals, 4 sauropsids (lizards and birds), 4 teleost fish and spotted gar. In addition, we also included 3R-ohnolog pairs and families in 4 teleost fish genomes. We used 5 non-vertebrate outgroups to identify 2R-ohnologs and 7 vertebrate outgroups to identify fish specific 3R-ohnologs (Figure 1 and Supplementary Table S1).

**Figure 1.**
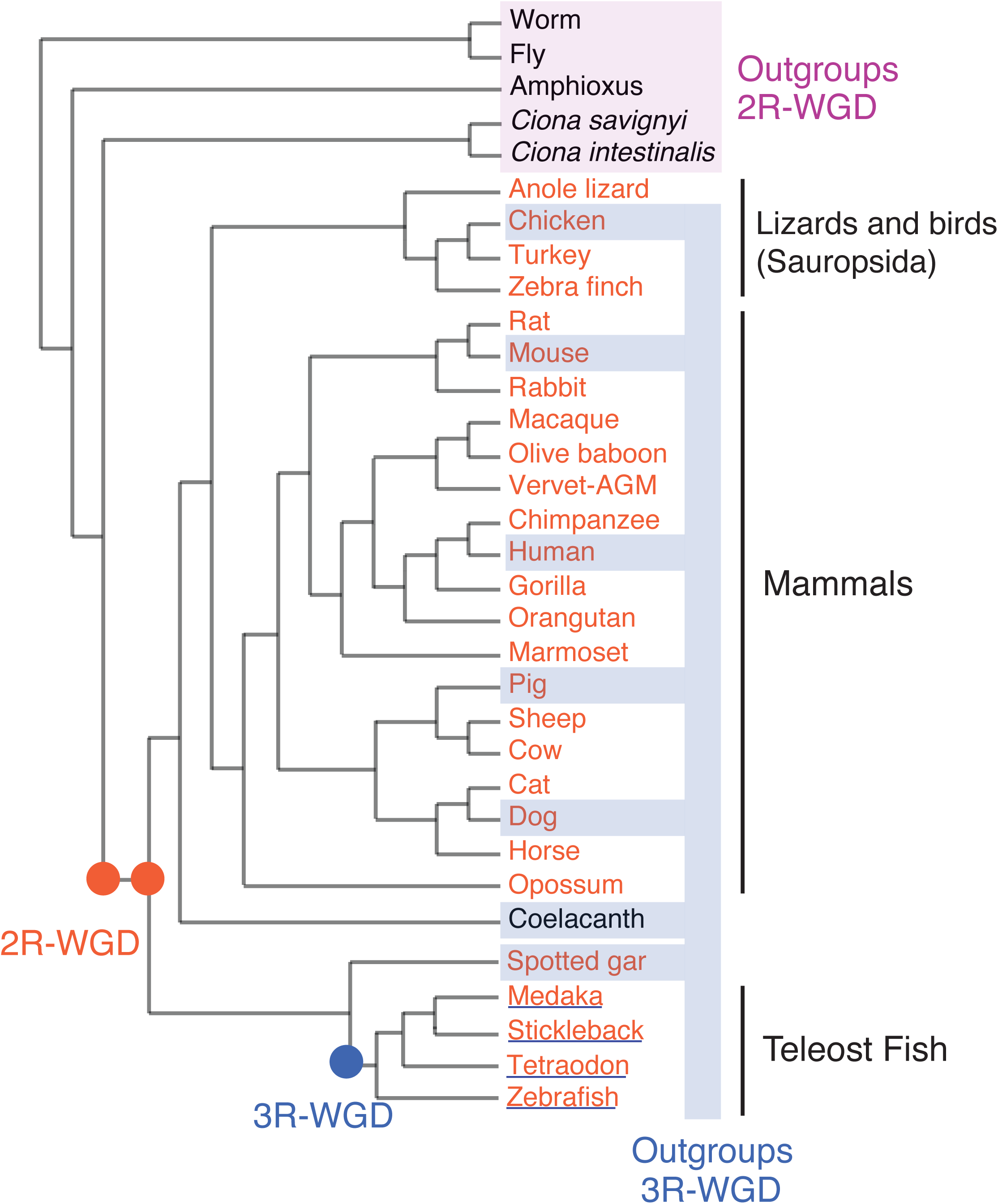
A schematic phylogeny (not scaled) of the organisms in the OHNOLOGS v2 database. Vertebrates analysed for 2R-WGD are in orange, and teleost fish species analysed for 3R-WGD are underlined. Outgroup species used to identify 2R- and 3R-ohnologs have been highlighted.

We collected genes (protein coding, micro-RNA, miscellaneous RNA, rRNA, snRNA, and snoRNA), their orthologs, paralogs and relative duplication node for all these organisms from Ensembl v84 using biomaRt (36–38). We chose these six classes of genes because they have information on orthologs and paralogs across many vertebrates, and the genes that had a lot of small-scale duplications (> 30) were excluded from analysis as they inflate the synteny calculations. Genome data for Amphioxus was obtained from JGI and amphioxus orthologs with other organisms were identified using BLASTp (8). To identify duplication time of paralogs consistently, we took the consensus timing across 7 Ensembl versions (v80 – v86).

We adapted the macro-synteny comparison approach, previously developed in (33), to identify ohnologs retained from both 2R-WGD (2R-ohnologs) and 3R-WGD (3R-ohnologs). Briefly, for each pair of outgroup and paleo-polyploid organisms, we first identified blocks of conserved macro-synteny using windows ranging from 100 to 500 genes (outgroup comparison). These macro-synteny blocks have a pattern of doubly-conserved synteny, where a window in the outgroup genome shares orthology with at least two other windows in the paleo-polyploid genome. The paralogs residing on these windows and duplicated at the time of 2R- or 3R-WGD are candidates for being 2R- or 3R-ohnologs, respectively. Similarly, we also identified syntenic windows by comparing each paleo-polyploid genome to itself (self comparison).

To refine these ohnologs further and eliminate spurious synteny patterns, we computed a quantitative score (called q-score) to assess the probability that any ohnolog pair could be identified by chance, following the approach developed in (33). In brief, all q-scores from different windows and outgroups were combined to give a global q-score for each ohnolog pair from outgroup comparison. Using multiple outgroups allowed us to identify ohnologs that may have moved to non-syntenic locations in some of the outgroup genomes. Similarly we obtained a q-score for self comparison to assess the chance of spurious association. In addition, while we used a simple geometric average of q-scores in (33), which cannot capture the gain of statistical power expected from the integration of multiple vertebrate genomes, here we developed a refined weighting scheme of species, which also takes into account the strong phylogenetically biased sampling of included species by using different weights for each vertebrate genome depending on its shared homology with other included genomes (see Supplementary Methods for details, Tables S2 & S3).

Using both self and outgroup averaged q-scores, we generated 3 sets of ohnologs (corresponding to strict, intermediate and relaxed criteria), and combined them into ohnolog families. Finally, we compiled both the 2R- and 3R-ohnolog pairs and ohnolog families for each organism in the interactive OHNOLOGS v2 database using Apache, CGI, Perl, Bootstrap and jQuery.

### Navigating the OHNOLOGS database

The home page lists all the organisms that are included in OHNOLOGS for 2R and 3R-WGD along with an introduction on ohnologs and WGDs. The search page allows a user to search for a gene symbol, Ensembl Id GO term or any keyword (Figure 2A). The search page also allows one to generate ohnolog families for any user-defined q-score criteria for a given organism. Upon a keyword or GO term query, all matching genes will be displayed along with their ohnolog status (Figure 2B). If a queried gene is an ohnolog, its ohnolog family will be displayed on the result page (for both 2R and 3R WGD for teleost fish) (Figure 2C and 2D). We show families for our strict q-score filter, and display the intermediate and relaxed families only if additional ohnologs are identified upon relaxing the q-score filter. The result page also includes links to pair page that has all ohnolog pairs that went into constructing that family (Figure 2E). The family result pages also links to the orthologous genes and ohnolog families in other vertebrates, to study the conservation of ohnolog families in other vertebrates.

**Figure 2.**
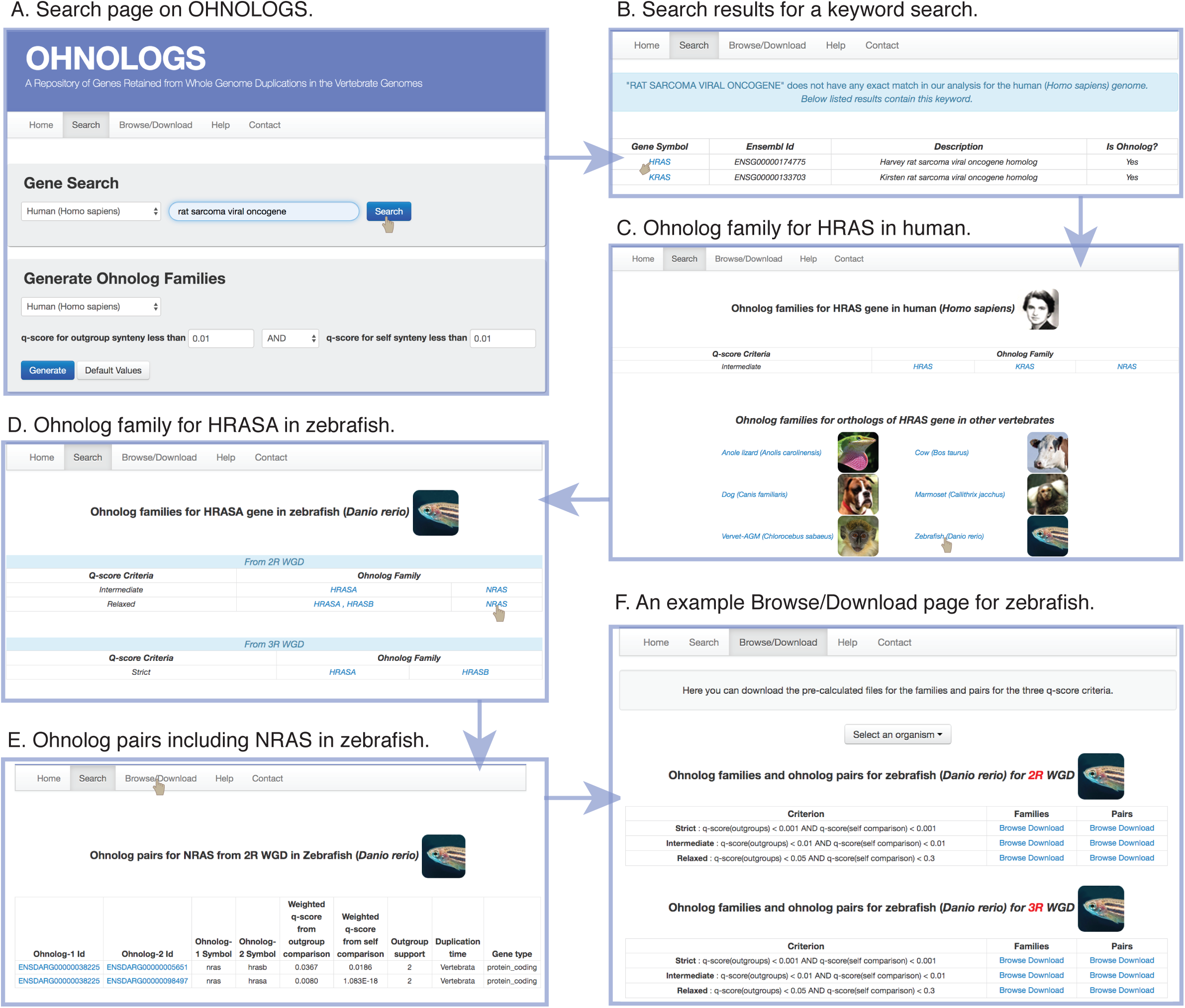
Navigating the OHNOLOGS database. A) Screenshot of the search page. B) Result page for a keyword search of “rat sarcoma viral oncogene” shows the matching genes in human. C) Ohnolog family page for HRAS gene in the human genome. D) From the family page, users can navigate to ortholog families in other vertebrates, e.g. zebrafish HRASA. E) Ohnolog pair page for zebrafish for NRAS gene. F) Browse/Download page for zebrafish showing both 2R and 3R-ohnolog pairs and families for all the three criteria.

The ohnolog pairs and families for our three pre-defined q-score filters can be explored and downloaded from the Browse/Download pages (Figure 2F). We link the genes on the browse pages to external databases including Ensembl, NCBI gene, GeneCards (for human), MGI (for mouse) and ZFIN (for zebrafish). The details of our approach, family descriptions and more details on q-score have also been included on the help page.

### Summary of the contents of the OHNOLOGS database

Using the expanded OHNOLOGS database we assessed the retention and loss of ohnologs across different vertebrates. We found that vertebrates have retained on average 25% 2R-ohnologs, which include two rounds of WGD, with teleost fish having subsequently retained on average 18% 3R-ohnologs (intermediate criteria) (Figure 3A, 3B and Supplementary Table S4). Sauropsids have retained the lowest numbers of 2R-ohnolog families, while teleost fish have retained the highest numbers of 2R-ohnolog families (Figure 3A). Yet, interestingly, we observe that more compact genomes, such as turkey and tetraodon, which typically contain also fewer genes, have retained about the same numbers of ohnologs than other birds or fish, respectively (Supplementary Table S4). This strong conservation of ohnologs across diverse genome sizes is consistent with their proposed retention mechanism through purifying selection in paleo-polyploid species (25,27,28).

**Figure 3.**
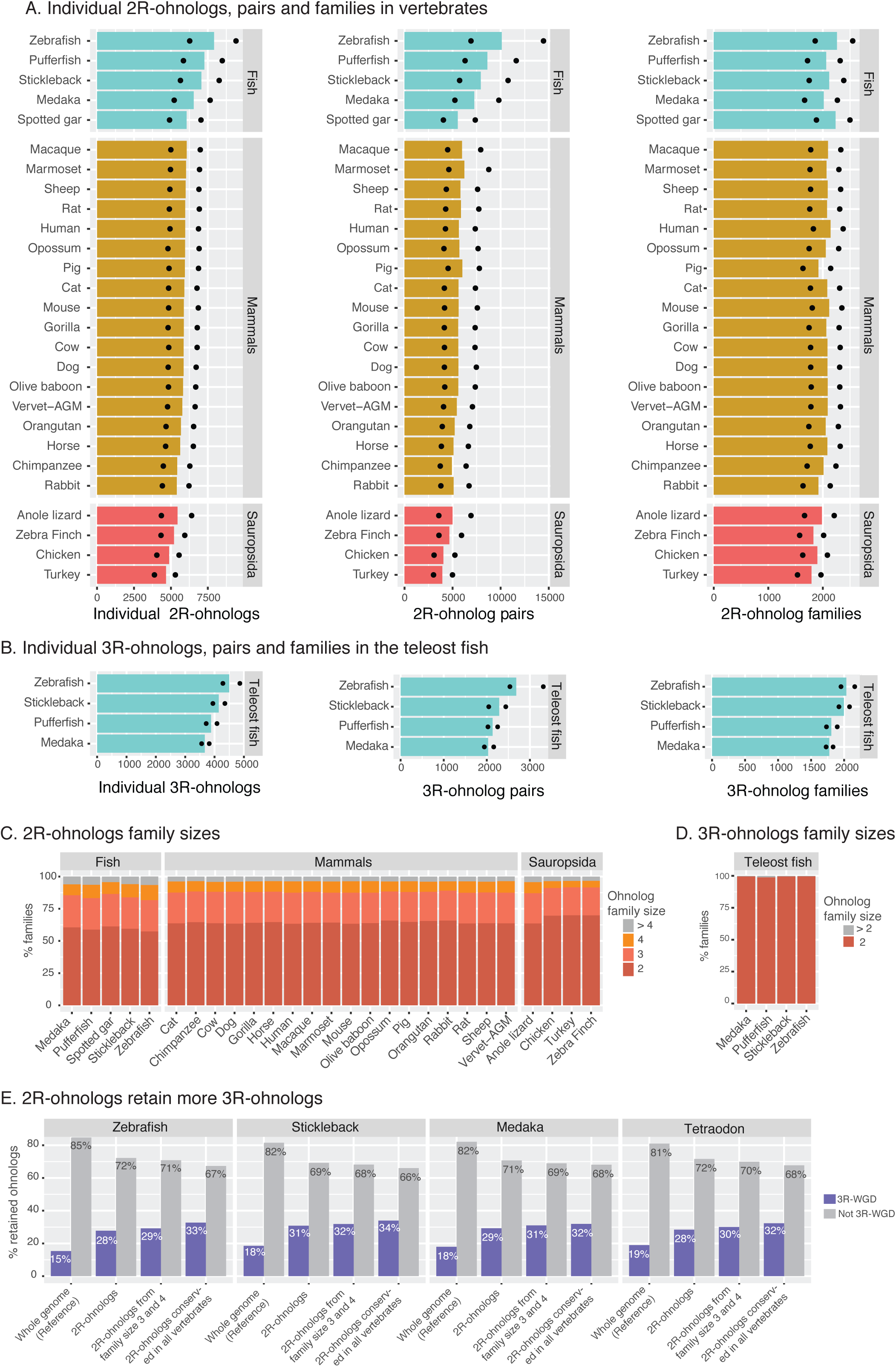
Description of the ohnolog genes, pairs and families in the database. A) Number of retained individual 2R-ohnolog genes, pairs and families in all the 27 vertebrates. Bars represent the numbers from the intermediate criteria. Ohnologs from strict and relaxed criteria are indicated by dots. B) Number of retained individual 3R-ohnolog genes, pairs and families in the 4 teleost fish species. Bars represent the numbers from the intermediate criteria. Ohnologs from strict and relaxed criteria are indicated by dots. C) Size of the 2R-ohnolog families from the intermediate criteria in vertebrates. Note that a vast majority of the families are of size 2, 3 or 4. D) Sizes of the 3R-ohnolog families from the intermediate criteria in the teleost fish hardly exceed size two. E) The 2R-ohnologs are significantly more likely to retain 3R-ohnologs, compared to genome-average. The retention of 3R-ohnologs is even higher for the 2R-ohnologs that belong to family size 3 or 4, and for the 2R-ohnologs conserved in all the 27 vertebrates. All the p-values are less than 1e-41, Chi-square test. Family counts are from the intermediate criteria.

A vast majority of retained ohnologs consists of protein-coding genes, while non-protein coding genes represent only a small fraction of ohnologs (Supplementary Table S5). For example, in human, out of the 7358 2R-ohnolog pairs from the relaxed criteria only 28 are mi-RNA ohnolog pairs and 2 are sno-RNA ohnolog pairs (Supplementary Table S5).

Remarkably, for all analysed vertebrates the size of 2R-ohnolog families rarely exceeds 4 ohnologs (Figure 3C), as expected for two rounds of WGD events. Similarly, virtually all 3R-ohnolog families are of size 2, as they are derived from just a single WGD event (Figure 3D). These family sizes also suggest a low rate of small-scale duplications and genome rearrangements following both 2R and 3R-WGD.

Next, the database can be used to analyse the branch-specific loss and retention of ohnologs. For instance, we found that 1,316 out of 2,373 ohnolog families with relaxed confidence criteria in human had an identical size in nearly all the 18 mammals (*i.e.* corresponding to a variance over mean size ratio lower than 0.1 across all 18 mammals). Then, out of these 1,316 conserved 2R-ohnolog families in mammals, 702 have the same size in teleost fish, including 396 families which also share the same size in sauropsids while the remaining 306 families correspond mainly to additional 2R-ohnolog losses in sauropsids; 119 families are larger in teleost fish and contain fish-specific 2R-ohnologs, while 86 families are smaller in teleost fish and correspond to 29 amniota-specific, 49 mammalia-specific and only 8 sauropsida-specific retentions of 2R-ohnologs.

Finally, we assessed if teleost fish with their additional 3R-WGD event tend to retain more ohnologs from the previous 2R-WGD events. Indeed, we found that in all four analysed teleost species, 2R-ohnologs tend to retain significantly more 3R-ohnologs (Figure 3E), in agreement with earlier reports (34). The retention of 3R-ohnologs is even higher for 2R-ohnologs that have retained 3 or 4 family members, and for the 2R-ohnologs that have been retained in all the 27 vertebrates (Figure 3E). For example zebrafish 2R-ohnologs from the intermediate criteria that have been also retained in all the analysed vertebrates are twice as likely to retain their 3R-ohnologs compared to genome-wide expectation (p = 5e-88, Chi-square test). This suggests that the evolutionary mechanism for the expansion of specific gene families through the retention of 2R-ohnologs (25,27,28) might also explain the biased retention of 3R-ohnologs.

## CONCLUSION

The updated OHNOLOGS v2 database is the most comprehensive resource available for the genes retained from both ancestral vertebrate 2R-WGDs and teleost fish specific 3R-WGD. It is based on a robust pipeline that downloads and processes datasets automatically using Ensembl, which makes it amenable to easy updates. We plan to expand and update OHNOLOGS periodically. Algorithmically, it is based on a quantitative comparative macro-synteny approach, which also takes into account the phylogenetically biased sampling of available vertebrate species. This approach assesses the confidence in each ohnolog pair and robustly identify ohnologs, despite lineage specific genome rearrangement, gene loss and small-scale duplication events. Using the datasets in OHNOLOGS we show a greater lineage specific ohnolog loss in sauropids compared to other vertebrate groups, and a high retention of 2R-ohnologs in subsequent 3R-WGD in teleost fish. In the light of the evolutionary significance of ancient WGDs and ohnologs for vertebrate evolution, the expanded and improved OHNOLOGS database should facilitate deeper comparative, evolutionary, genomic and functional analyses of the ohnolog genes in vertebrates.

## AVAILABILITY

All the data and code used to construct OHNOLOGS is available at http://ohnologs.curie.fr and its associated GitHub repository at https://github.com/param-p-singh/Ohnologs-v2.0.

## SUPPLEMENTARY DATA

Supplementary Data are available at NAR online.

## ACKNOWLEDGEMENT

We acknowledge technical support from the service informatique of Institut Curie for hosting and maintaining the server infrastructure. We thank Vincent Cabeli and Marcel Ribeiro Dantas for help in updating the server.

## Supplementary Methods

### 1 Confidence assessment of ohnolog pairs: combining q-scores

#### 1.1 Ohnologs v1 database: a simple average of log q-scores over vertebrate species

The confidence assessment of individual ohnolog pairs in the original Ohnologs v1 database (*1*) relied on the definition of quantitative outgroup and self-synteny scores (q-scores) for each vertebrate species, *i.e.*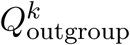 and 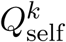, where *k* =human, mouse, rat, pig, dog and chicken. See Singh *et al.* PLoS Comput Biol 2015 paper (*1*) for a detailed computation of q-scores from synteny comparison.

Then, to circumvent the difficulty to identify ohnolog pairs in each vertebrate genome due to lineage specific rearrangement, gene loss and small scale duplication events, geometric averages of outgroup and self-synteny q-scores were taken over the six amniote species included in the Ohnologs v1 database:

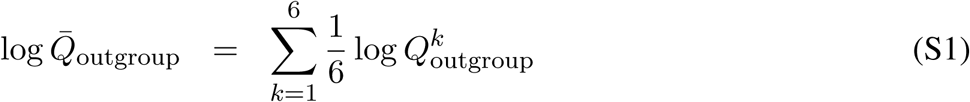

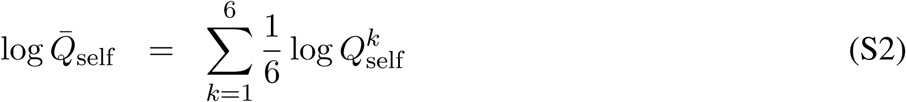

Using such averaged q-scores was shown to improve the statistical significance of the inferred ohnologs by allowing to identify ohnolog pairs that are no longer in significant synteny in a particular vertebrate genome, if their respective orthologs form a high confidence ohnolog pair in other vertebrates.

However, simple q-score averages fall short of (i) assessing the gain in statistical power expected from the integration of multiple vertebrate species (as the weights 1/6 in Eqs. S1 and S2 sum to 1), as well as, (ii) taking into account the phylogenetically biased sampling of vertebrate species by using equal weights (1/6), while the more recently diverged mouse and rat genomes are expected to bring rather redundant information on ohnolog retention as compared to phylogenetically more distant species such as chicken.

#### 1.2 Ohnologs v2 database: a weighted sum of log q-scores over vertebrate species

The expanded Ohnologs v2 database addresses the shortcomings on the statistical confidence of ohnolog pairs from the original Ohnologs v1 database.

To this end, we modified the definitions of outgroup and self-synteny q-scores, from Eqs. S1 and S2, as weighted sums of log q-scores over all *N* = 27 vertebrate species included in the Ohnologs v2 database for 2R-ohnologs and all *N* = 4 included teleost fish species for 3R-ohnologs (see Table S1),

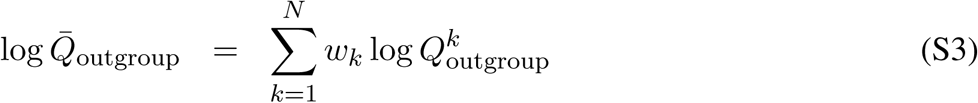

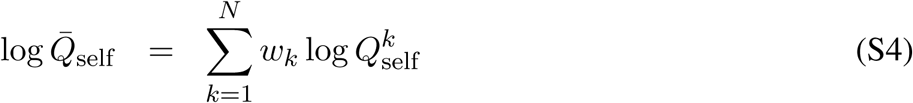

where the weights (*w*_*k*_) are meant to (i) capture the gain in statistical power expected from the integration of 27 vertebrates including 4 teleost fish species (*i.e.* Σ_*k*_ *w*_*k*_ > 1) and (ii) take into account the strong phylogenetically biased sampling of included species by using different weights for each vertebrate genome depending on its shared homology with other included genomes.

The computation of the individual weights, 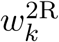 for 2R-ohnologs and 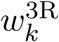 for 3R-ohnologs, are detailed in the following section. It is based on the times of divergence between each pairof vertebrate genomes included in the study (Table S2) and the values of 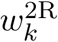 and 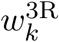 are listed in Table S3.

The overall gain of statistical power is estimated as 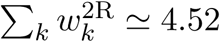 for 2R-ohnologs and 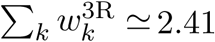 for 3R-ohnologs. This corresponds to an effective number of “independent species” of about 4.5 out of the 27 included vertebrates for assessing the confidence of 2R-ohnologs and to an effective number of “independent species” of about 2.4 out of the 4 included teleost fish for assessing the confidence of 3R-ohnologs.

In addition, as anticipated, recently diverged species of overrepresented vertebrate subgroups are assigned very small weights, which only amount to a very small fraction of the total weight. In particular, each of the 8 included primates has an individual weight around 0.01-0.02, while the sole representatives of long diverged subgroups have proportionally very large weights, such as Spotted Gar (*w* ≃ 0.72) or Anole Lizard (*w* ≃ 0.57).

A consequence of the gain of statistical power between Ohnologs v1 and v2 databases is that we could define more stringent confidence criteria for ohnolog pairs and generated ohnolog families as,

- strict 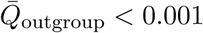 AND 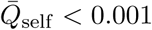
- intermediate 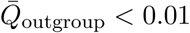 AND 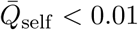
- relax 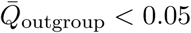 AND 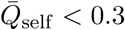

### 2 Weighting scheme for phylogenetically related sequences

As discussed above, the effective number *N*′ of “independent species” is smaller than the actual number *N* of phylogenetically related species included in the analysis.

One way to estimate *N*′ and the corresponding weights *w*_*k*_ for each phylogenetically related species (with 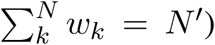) is through the apparent increase of variance of an ordinal character *x* (such as the number of genome rearrangements) across *N* non-independent species. The result is quite general and the increase of variance can be used to infer consistent weights for a generic dataset of *N* non-independent samples, as discussed in the next section.

#### 2.1 Generic increase of variance due to non-independent samples

The generic increase of variance between the *N* non-independent samples can be illustrated on the example of a theoretical dataset with *N*′ independent samples, each repeated *n*_*k*′_ times (or not repeated if *n*_*k′*_ = 1) to yield a larger dataset of 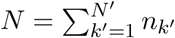 non-independent samples.

The variance obtained for the larger non-independent dataset of size *N* reads:

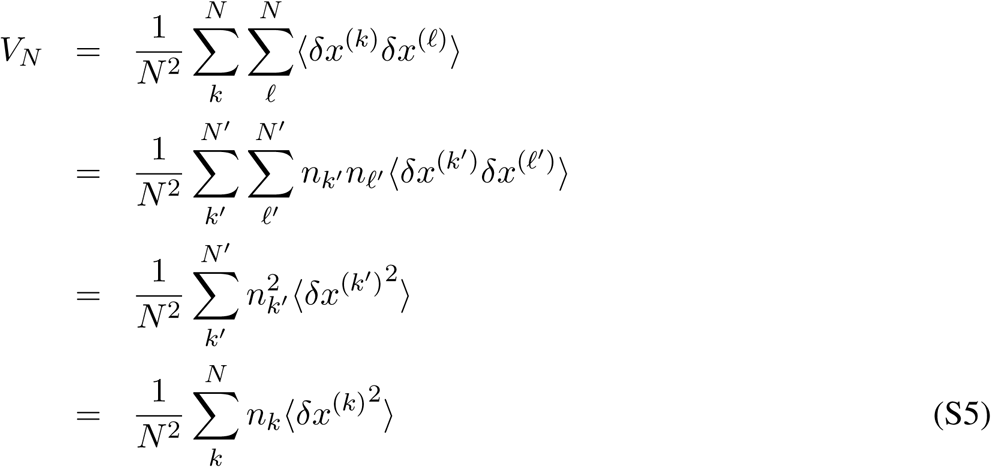

as 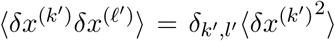 for independent samples and using 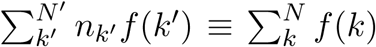 with *n*_*k*_ = *n*_*k′*_ for each of the *n*_*k′*_ samples *k* that are duplicates of sample *k*′.

When all samples are independent, that is, if *n*_*k*_ = 1 for all *N* samples, one recovers the well known results (adopting the rescaling 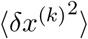 = 1 for all *k*),

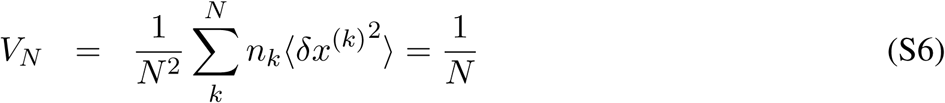

By contrast when the samples are not all independent, that is, if *n*_*k*_ > 1 for some of the *N* samples 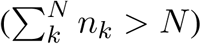, one gets

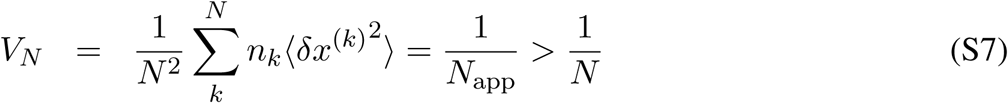

as if the apparent number of independent samples was smaller, *N*_app_ < *N*.

This suggests to weight each non-independent sample *k* with a probability weight, *w*_*k*_ = 1*/n*_*k*_ ⩽ 1 with 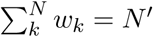 and to define the corrected variance for effective sample size as,

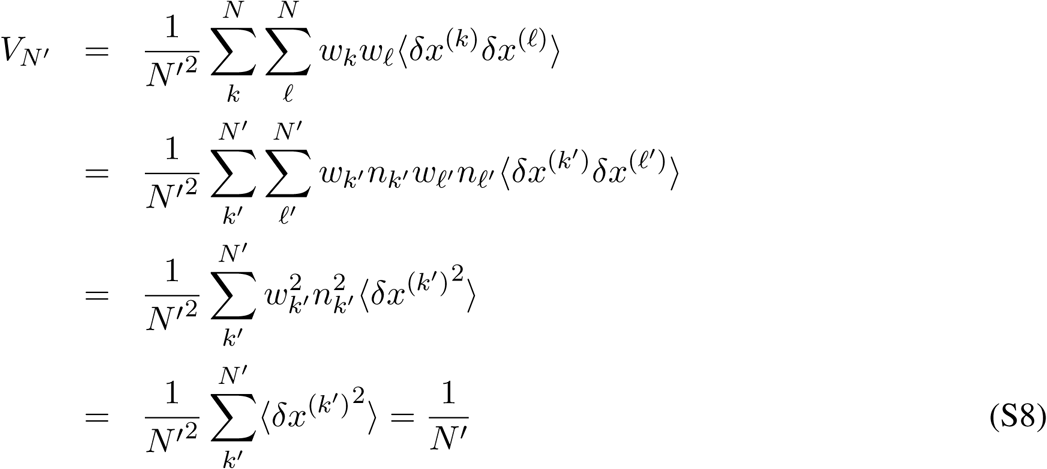

using *w*_*k*_*n*_*k*_ = 1. Note that *V*_*N*′_ can also be expressed in the actual sample space with *N* non-independent samples, instead of the effective sample space with *N*′ independent samples (which are not typically known), as,

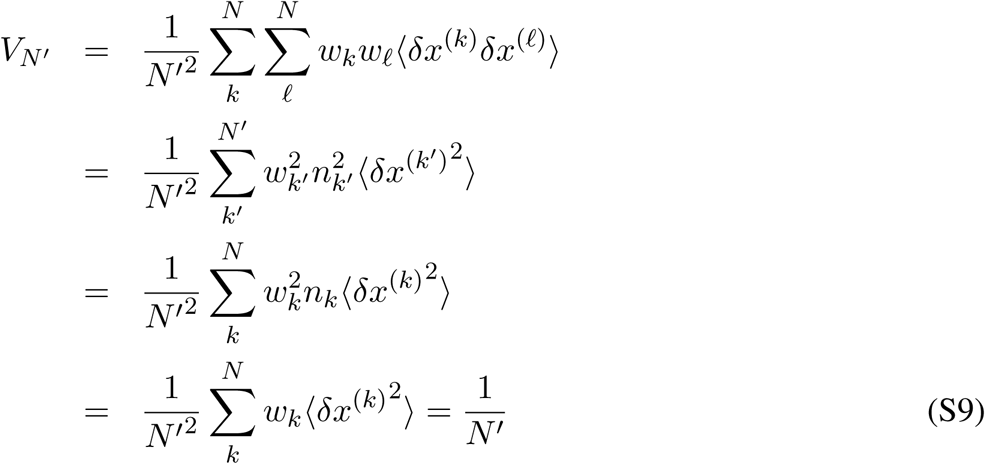

using 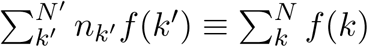 and ∀*k, w*_*k*_*n*_*k*_ = 1 and 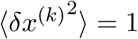.

#### 2.2 Sample weighting scheme by inversion of the variance-covariance matrix

The above results show that the sample weights {*w*_*k*_} are solutions of the following equation,

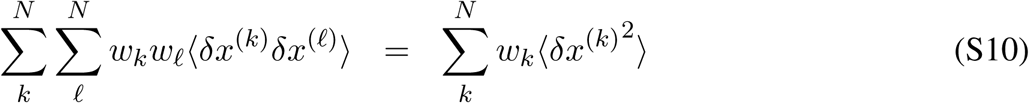

While Eq. S10 with *N* unknown weights is underdetermined, one can easily show that this equation also applies to individual summand for each *k* as,

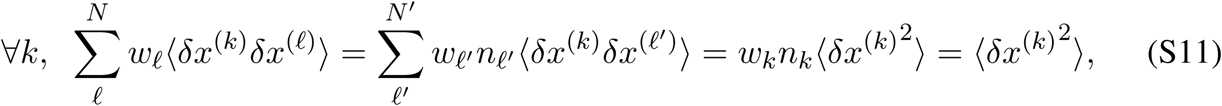

using 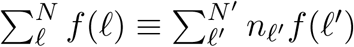 and ∀*k, w*_*k*_*n*_*k*_ = 1.

Eq. S11 can be written in the following matrix form, after rescaling *δx*^(*k*)^ by its mean deviation as 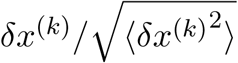,

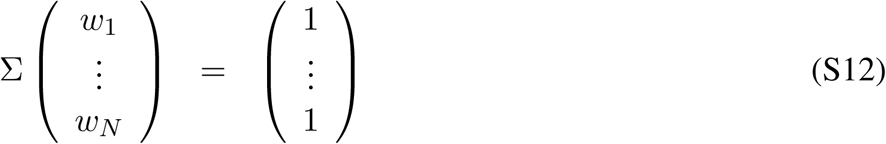

where Σ = [Σ_*kℓ*_] with 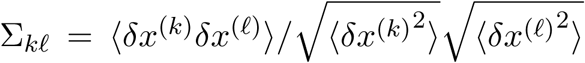 is the rescaled variance-covariance matrix between samples, which leads to the following weight solution whenever the variance-covariance matrix is invertible,

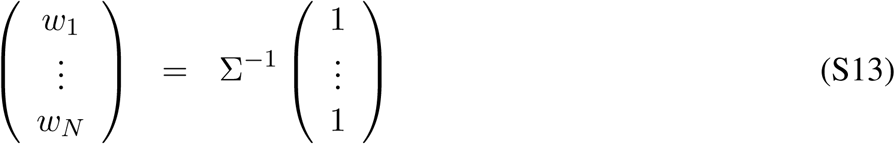

While Eq. S13 seems to give the straightforward solution of the generic sample weighting problem, in practice, the variance-covariance matrix Σ is typically not invertible. In particular, straightforward averages of variances and covariances over the available samples, which imply 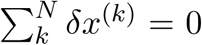, yields a singular variance-covariance matrix (as all rows and columns sum to zero).

Yet, in some particular cases, the form of the variance-covariance matrix can be conjectured independently from the specific data of interest and used to solve Eq. S13.

This is the case for time series of dynamical systems with exponential relaxation over time (*2, 3*) or for phylogenetically related sequences (*4, 5*), as discussed in the next section.

#### 2.3 Application to weighting phylogenetically related sequences

The form adopted for the variance-covariance matrix of phylogenetically related sequences is directly inspired by the form proposed by Altschul *et al* in ref. (*5*) to estimate weights of sequence data related by a tree.

Following these authors, we reason that as genome rearrangements and gene loss events accumulated in ancestral vertebrate genomes after each WGD event, the distance of their alignment with the reference paleoploid genome progressively shift. At first approximation, one expect a *linear* accumulation of some finite number (*X*) of genome rearrangements and gene loss events over time, as these evolutionary changes are essentially non-reversible (small scale duplication events might even lead to exponential growth of gene families over time (*6*)). This is to be contrasted with an unbiased reversible random walk in sequence space, which would lead to a purely diffusive dynamics with a sublinear (square root) accumulations of changes over time.

Hence, due to this progressive shift in genome space, the variance of accumulated changes, *X*_*k*_, of a given vertebrate genome, *G*_*k*_, is expected to increase *quadratically* with time *t*_*k*_ since a WGD event, *i.e.* 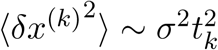, instead of linearly with time for a perfectly diffusive dynamics.

Similarly, the covariance of accumulated changes in two genomes, *G*_*k*_ and *G*_*ℓ*_, having diverged after some time *t*_*kℓ*_ after a WGD event is expected to increase *quadratically* as, 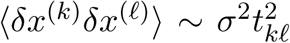, assuming that subsequent changes after the two genomes have diverged were completely independent and could not therefore further increase the covariance.

All in all, this leads to the following form for the rescaled variance-covariance matrix, Σ = [Σ_*kℓ*_], where 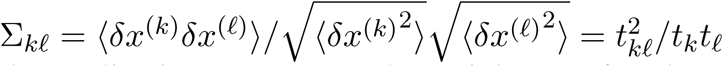, that is independent for the prefactor *σ*^2^.

In the application to compute the weight *w*_*k*_ of each species, the times of 2R-WGD and 3R-WGD were estimated by averaging recent estimates as *t*_2*R*_ = 535 MY (*7–10*) and *t*_3*R*_ = 328 MY (*11–15*), respectively, and the times since divergence of each pairs of species, *d*_*kℓ*_ = *t*_*WGD*_ − *t*_*kℓ*_, were taken from the TimeTree database (*16*) and listed in Table S2. The final values for 2R- and 3R-WGD weights are listed in Table S3.

